# The rotating tilted lines illusion for the evaluation of cognitive abnormalities

**DOI:** 10.64898/2026.03.05.709956

**Authors:** Chris W. Bao, Emma Martin, Basilis Zikopoulos, Arash Yazdanbakhsh

## Abstract

**Background:** The population receptive field (pRF) in vision reflects the functional receptive field arising from millions of overlapping single receptive fields across visual areas and eccentricities. pRFs are typically estimated with fMRI to gain insight into visual processing. Alternative methods of pRF estimation, such as using optical illusions, have been explored only sparingly. In this study, we explore the rotating tilted lines illusion (RTLI), in which a circle formed by tilted lines appears to rotate as it expands or contracts in the visual field (e.g., from moving the head back and forth).

**New Method:** We propose a novel set of computer-generated animations of the RTLI that measure the visual and temporal characteristics of the illusory rotation, enabling quantitative estimation of the spatial extent and temporal dynamics of the pRF.

**Results:** We derived pRF size estimates consistent with those estimated from fMRI and electrophysiological methods. We then projected changes in RTLI percept trends according to abnormalities in visual processing in autism spectrum disorder (ASD), schizophrenia (SZ), aging, and Alzheimer’s disease (AD).

**Comparison with existing methods:** Compared to fMRI and electrophysiology, RTLI-based pRF estimation is accessible, low-cost, and feasible at home or during inpatient visits without specialized equipment.

**Conclusions:** We show that our novel method can approximate pRFs, which in turn can be potentially applied for early detection, probing the progress, and treatment screening in AD, SZ and ASD.

## 1. Introduction

For over a century, modern psychophysics has embraced optical illusions as a powerful research tool in probing human perception. By leveraging disparities between physical visual input and perceptual experience, illusions unravel the mechanisms and biases of the human visual system and pathways. For example, the Kanizsa triangle and square reveal our tendency to see shapes that do not actually exist in full, while apparent motion reveals how the visual system infers motion from distinct, transient stimuli that are spatially and temporally separated (Kanizsa, 1976; Layton et al., 2012; Neumann et al., 2007; Nishina et al., 2007; Wertheimer, 1912; Wurbs et al., 2013; Yazdanbakhsh and Watanabe, 2004; Zhu et al., 2023). Illusions, more recently, have also been utilized to study perceptual aberrations in disorders (Klein et al., 2020; Montiel et al., 2025; Sperandio et al., 2023). Motion illusions in particular have been considered alongside other tools to probe neurodevelopmental disorders such as developmental dyslexia (DD) and autism spectrum disorder (ASD) (Gori et al., 2016).

The rotating tilted lines illusion (RTLI) is a motion illusion in which a circle formed by tilted lines appears to rotate as it expands or contracts in the visual field (e.g., by moving the head back and forth). The viewer infers such illusory motion because the RTLI capitalizes on a well-known phenomenon in vision science known as the aperture problem (see section 2, theory and calculation). The RTLI, by triggering illusory rotation and potentially engaging the magnocellular pathway, can be a useful visual illusion candidate to be utilized in the detection of DD in children (Gori et al., 2016, 2014) by estimating the population receptive field size (pRF) (Yazdanbakhsh & Gori, 2008). pRF size is one of the key metrics for probing neurodevelopmental, neuropsychiatric, and neurodegenerative disorders (Elul and Levin, 2024). As a cost-effective method of measuring pRF sizes, the proposed RTLI task is an alternative and accessible method that can contribute in assessing disorders associated with deviations in pRF size, such as autism spectrum disorder (ASD), schizophrenia (SZ), aging and Alzheimer’s disease (AD). Based on our prior work on computational analysis of RTLI (Yazdanbakhsh & Gori, 2008), and empirical data collection with full RTLI spatial and temporal parameters, in this study we a) fully characterized the RTLI percept in fine grained spatial and temporal configurations for neurotypical human observers, and then b) predicted RTLI percept outcomes for ASD, SZ, aging, and AD, in the light of published data about the pRF changes in associated clinical populations.

## 2. Theory and Calculation

### 2.1 The aperture problem

When neurons in the visual cortex process motion, their “sight” does not encompass the entire visual field. Instead, their *receptive field* (RF) is a limited area, akin to a viewing window or aperture. This condition creates the “aperture problem” when evaluating the motion of sufficiently long contours or lines. Figure 1 shows sample stimulus configurations that either generate or avoid the aperture problem. When a receptive field (RF) is larger than the contour, the entire line is visible, and the neuron’s response aligns with the true motion (Figure 1A). When the RF is smaller than the contour but still includes at least one of the line’s endpoints (terminators), the neuron can still represent true motion through extrapolation (Figure 1B). The aperture problem arises when the RF is smaller than the contour and excludes its endpoints, as shown in Figure 1C. The visual signal is ambiguous; with no anchoring reference point to track the motion, the true velocity may lie anywhere in a 180° arc on the side of contour progressing displacement (Figure 1C). In this case, only a motion component which is orthogonal to the line is detected. The barber pole serves as a familiar example: instead of perceiving a turning cylinder, the viewer sees diagonal lines moving vertically which again demonstrates that the visual system relies on local motion cues, similar in Fig.1 A-B to infer the overall motion, leading to misinterpretations. The barber pole motion is horizontal, but viewed through a narrow, vertical aperture, the visual system perceives the motion vertically, because the motion of the grating’s edges along the vertical sides of the aperture is the most prominent cue.

**Figure 1.**
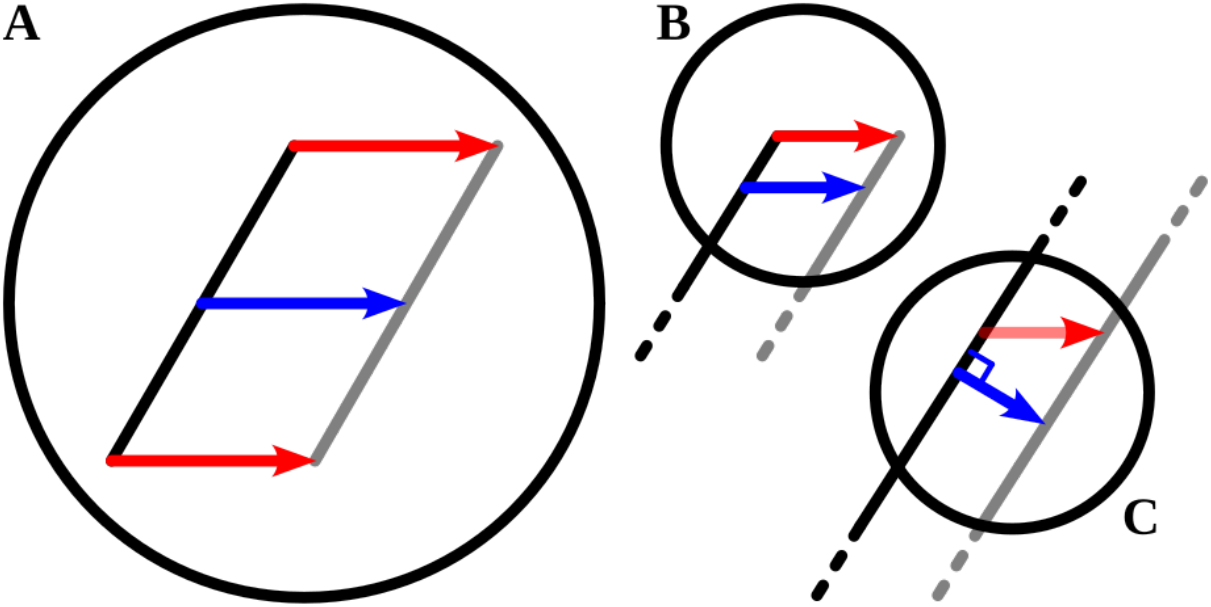
Receptive fields and the aperture problem. Case A: receptive field (black circle) encloses the moving line; true motion can be detected. Case B: receptive field encloses an endpoint; true motion can be detected and extrapolated from endpoint motion. Case C: receptive field within the moving line; aperture problem takes effect, only perpendicular to the line component of motion can be detected. Red indicates true motion while blue indicates perceived motion.

### 2.2 Illusory motion perception in the rotating-tilted-lines-illusion

The RTLI is a circle composed of lines tilted with respect to the radius at each circumference point (Figure 2A). Illusory motion appears when the stimulus expands and contracts in the visual field, which can be achieved either by moving the head back and forth or directly demonstrating expanding and contracting the stimulus on a computer monitor. Figure 2B demonstrates the aperture problem in the context of the RTLI: the line segment extends beyond the sample RF, causing perceived motion to appear orthogonal to the line segment, as in Figure 1C. This perceived motion differs from the true motion produced by expansion and contraction of the RTLI, creating a tangential motion component that gives rise to the appearance of rotation.

**Figure 2.**
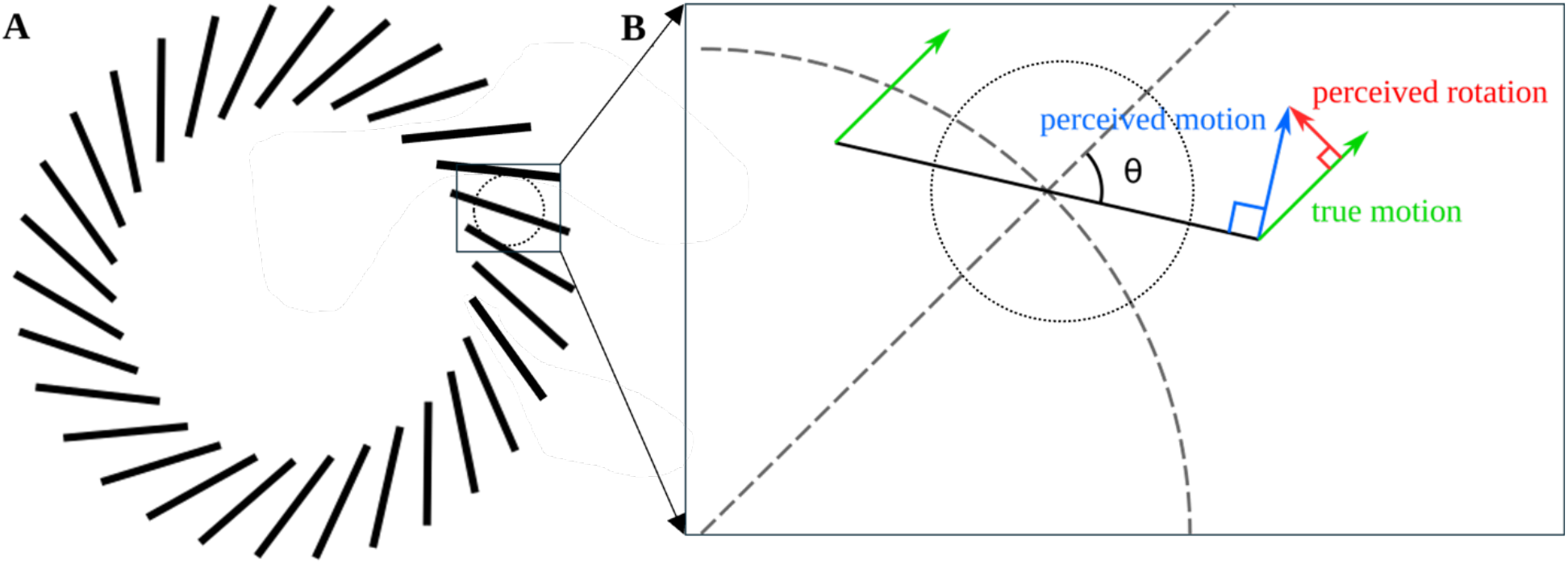
Illusory motion in the rotating tilted lines illusion. A: The rotating tilted lines illusion. Sample RF is represented by the dotted circle. **B:** RTLI, zoomed in to show one line segment with sample RF. True motion (green) arises from the RTLI expanding and contracting in the visual field. The aperture problem creates perceived motion (blue) which is orthogonal to stimulus line, and perceived rotation (red) as a component.

### 2.3 Estimating population receptive field size using the RTLI

The extent of illusory rotation in the RTLI can be modified by varying stimulus parameters. For instance, if the line segment in Figure 2B were shortened to fit entirely within the RF, the perceived and true motion vectors would align, eliminating the rotational illusion. Systematically varying line length and measuring the transition from rotational to non-rotational percepts thus allows for estimation of RF size. However, unlike the scenario depicted in Fig. 2B, the RTLI task does not probe individual RFs and rather probes population average of many overlapping receptive fields. Across and within visual areas, regions, and eccentricities, receptive fields have a range of scales and sizes; receptive field size increases with eccentricity and along a visual hierarchy (Bakin et al., 2000; de Best et al., 2019; Layton et al., 2012; Oliver W. Layton et al., 2014; Layton and Yazdanbakhsh, 2015; Qian and Yazdanbakhsh, 2015; Wurbs et al., 2013). Millions of overlapping receptive fields together support visual perception across the entire visual field (Layton et al., 2014). The RTLI task estimates the size of the *population* receptive field (pRF), the functional receptive field resulting from motion-processing units in the visual system. The magnitude of illusory rotation reflects the proportion of receptive fields (RFs) affected by the aperture problem. In simultaneously sampling all size and position scales (i.e., the population of RFs), this psychophysical method captures the heterogeneity of RF sizes across the cortex. By systematically varying the length of RTLI lines, the heterogeneity can be quantified: shorter lines engage fewer RFs subject to the aperture problem, while longer lines engage more. When illusion strength reaches its maximum, indicating that nearly *all* RFs are affected by the aperture problem, further increases in line length produce little to no change. Therefore, all tested line lengths including those at which illusion strength begins to plateau quantify RF heterogeneity and provide an estimate of pRF size.

## 3. Methods

### 3.1 Participants

16 participants were recruited and tested from the Boston University undergraduate student pool (age range 18–22). All procedures were approved by the Boston University Charles River Campus Institutional Review Board (3651E), and written consent was obtained according to the social and behavioral focused human subjects protection. All participants had normal or corrected-to-normal vision.

### 3.2 Stimuli

Stimuli were generated with the Tkinter module in Python 3.7 and displayed on an Eizo Corporation ColorEdge CG241W 24-inch (518.4 × 324.0 mm) LCD monitor. The resolution was 1920 × 1080 pixels with a 60 Hz refresh rate. A chinrest was used to control the distance between the eyes and the monitor, *d* ≈ 43 cm.

Each stimulus was made up of 60-line segments arranged in a circle and angled relative to the radial line (Figure 3). Three parameters of the illusion were varied in this study: line length, stimulus radius, and animation speed. Line angle was set at 45°, as our pilot studies showed that the optimal illusion strength was achieved at angles between 40–60° from the radial line, as predicted by the aperture problem geometry (Appendix B). Maximum displacement, or amplitude of the contraction/dilation, was set at 100 pixels (3.6° visual angle) from the starting position to each side. Whereas Yazdanbakhsh et al. (2008) scaled displacement with line length, the present study used a fixed displacement across all conditions to control for velocity. Varying displacement with line length would otherwise increase line velocity and potentially exaggerate illusory motion strength. Moreover, the animations were not strictly scaling, as if the illusion would be if the head was moved back and forth; instead, the line lengths were kept constant while the segments moved radially. This was done for simplicity and to reduce errors resulting from scaling widths up and down. A cross (plus sign) was displayed at the center of the screen for fixation (Figure 4). Viewing was binocular.

**Figure 3.**
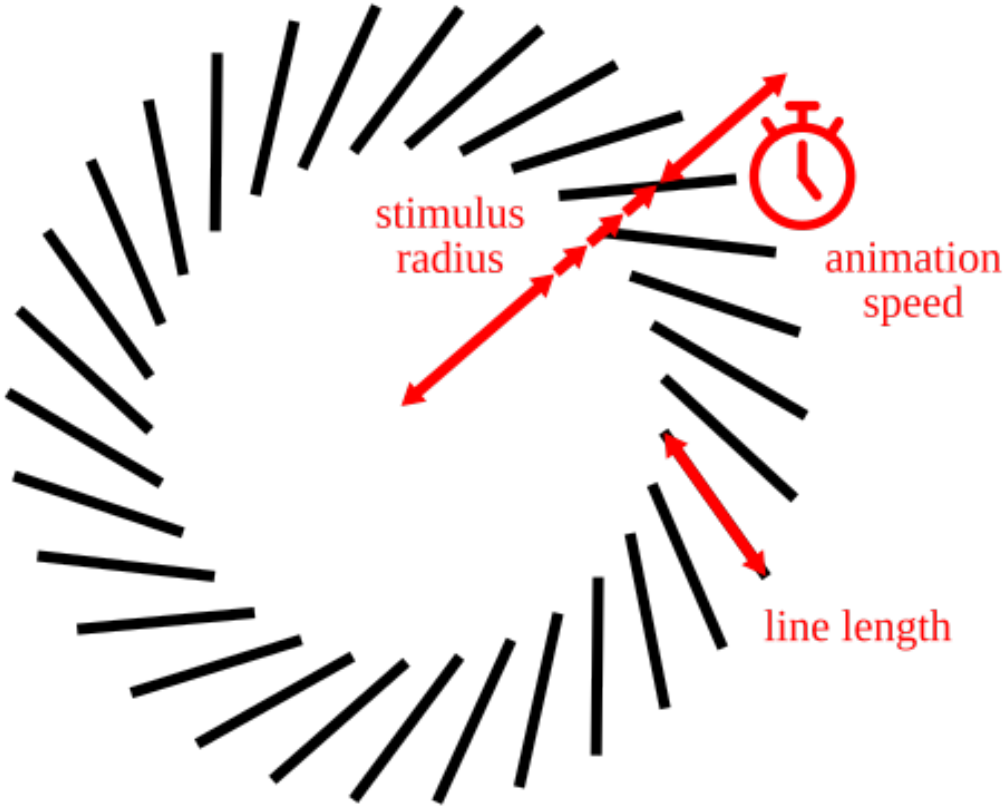
The rotating tilted lines illusion. This experiment studied the effects of three parameters on illusion strength: animation speed, stimulus radius, and line length.

**Figure 4.**
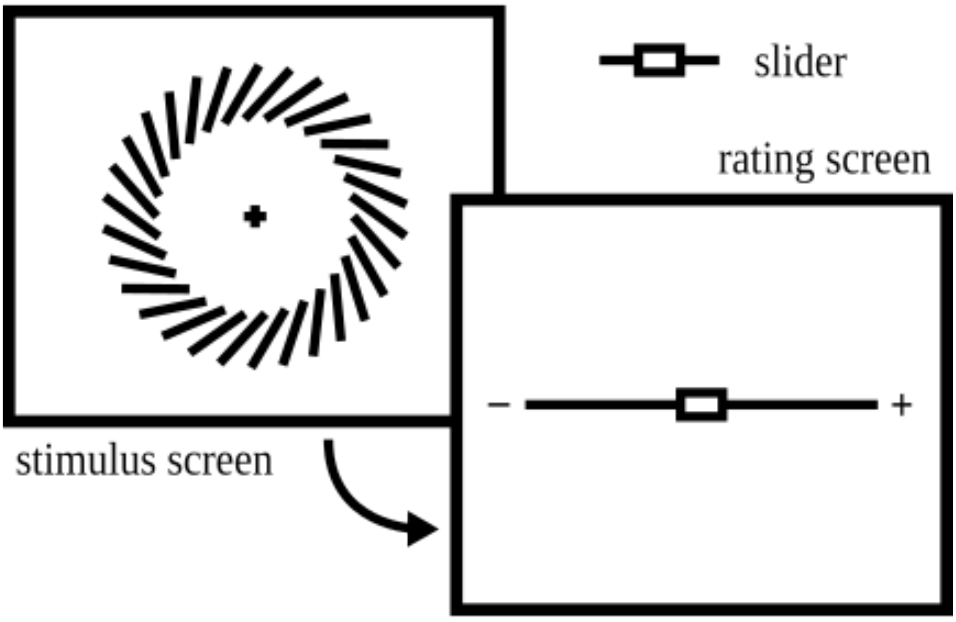
The stimulus and rating screens. Each block alternated between presentations of the illusion and sliders to rank the relative illusion strength.

### 3.3 Procedure

Because each of the three variables (line length, stimulus radius, and animation speed) have five levels, the experimental design was 5^3^ factorial, producing 125 total combinations. A block design was used, wherein each block included one of every combination. After being instructed in the use of the rating slider, buttons to proceed and exit, and background information on the illusion, subjects were presented with one practice block and three experimental blocks with rest periods in between. Ratings from the practice block were not recorded, allowing the subjects to familiarize themselves with the illusions and their relative strengths. As the scale is subjective and relative, they would need to have seen the full range of strengths before recorded results could be accurate. The procedure length including all blocks lasted from 60 to 90 minutes, depending on how quickly the observer proceeded through the task.

During each trial, the subject was presented with a three-second-long looping (expanding-contracting) animation of the illusion. This duration was selected to accommodate sufficient viewing time while preventing excessive fatigue in subjects given the sheer volume of combinations. Subjects were then instructed to rate the illusion power with a slider on the screen (Figure 4). The position on the far left corresponded to 0 or no rotation, and the position on the far right corresponded to 100 or the strongest rotation they had seen. A slider was used in place of, e.g., a Likert-type scale for greater continuity and granularity. Explicit numbering was removed so as to remove bias for certain numbers, such as even numbers or multiples of five or ten. The slider was reset to zero (midpoint) between each trial in an effort to make the ratings as independent from previous trials as possible.

### 3.4 Data Analysis

The data were analyzed with the Python numpy v1.23.0, scipy v1.9.0 and statsmodels v0.13.2 packages. Data were pooled across subjects. For analyses across one or two main variables, data were pooled across the rest. For example, when examining illusion strength vs. line length, data across all levels of stimulus radius and animation speed were pooled together.

## 4. Results

### 4.1 Effects of RTLI parameters on illusion strength

Three 2-way ANOVA tests were performed to identify relationships among each pair of explanatory variables. Interactions were found between line length and stimulus radius (*F*(16) = 4.547, *p* < 0.05) and between line length and animation period (*F*(16) = 1.714, *p* < 0.05). Interactions between stimulus radius and animation period were not significant (*F*(16) = 0.444, *p* > 0.05). Kruskal-Wallis tests revealed significant associations from all three main variables to illusion strength (*H* = 4705, 9233, 4143, all *p* < 0.05). Each variable affected the illusion strength in a different manner. Increasing line length monotonically increased the strength of the illusion (Figure 5A). Very short line segments (≤ 0.696°) created no significant rotation, but longer line lengths (≥ 2.09°) produced diminishing increases (seen as a decreasing slope in the graph). Conversely, increasing stimulus radius monotonically decreased the strength of the illusion (Figure 5B). Animation period, a discrete variable ranging from 0.25 to 2.0 seconds, had the highest variability in illusion strength across trials. The trend was that increasing the period (i.e., decreasing animation speed) also decreases illusion strength (Figure 5C).

**Figure 5.**
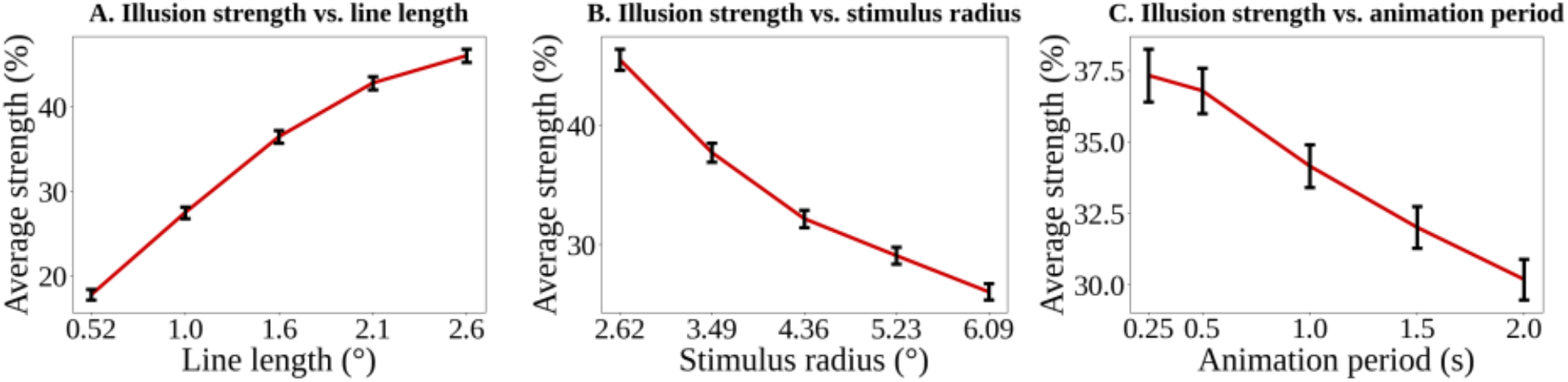
Effects of line length, stimulus radius, and animation period on illusion strength. A: illusion strength monotonically increases with line length. B: illusion strength monotonically decreases with stimulus radius (distance of lines from center of vision). C: illusion strength has a trend of decreasing with animation period (increases with animation speed). Error bars show 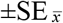. “°” symbol denotes degrees of visual angle. See Appendix C for more detailed plots.

Pearson, Spearman, and Kendall correlations were computed for the relationships between each parameter and illusion strength. All results were significant, with line length having the strongest correlation (*r* = 0.37, *ρ* = 0.39, *τ* = 0.29) and having *p*-values of approximately 10^-200^ for all three tests. Stimulus radius had an intermediate effect (*r* = –0.24, *ρ* = –0.23, *τ* = –0.17) with *p*-values of about 10^-75^. As expected from the higher variance, animation period had the weakest, although still significant, correlation with illusion strength (*r* = –0.10, *ρ* = –0.07, *τ* = –0.05, *p* ≈ 10^-10^).

### 4.2 pRF estimation

Using the length at which illusion strength started to plateau as an estimate of the maximum pRF size (Yazdanbakhsh and Gori, 2008a), we obtained values of between 1.56 and 2.09°. These bounds are in line with V3 to middle-temporal RF sizes given in fMRI and electrophysiological literature. For example, Dumoulin & Wandell (2008) obtained fMRI estimates of 1.2 to 2.4° for V3 RFs at the ranges we tested. Albright & Desimone (1987) used a single-neuron recording procedure to obtain a linear model predicting sizes about 2.5 to 5° in the middle-temporal region of macaques.

Figure 5A, which plots illusion strength against line length, can further be used to deduce the size distribution of receptive fields processing the motion. To reiterate: as the lines in the illusion increase in length, they will exceed the bounds of RFs and cause the aperture problem; the ratio of these ‘fooled’ RFs to the total is the strength of the illusion (see section 2.3). If one continues till taking the highest data point –maximal illusory strength– to have fooled 100 % of the RFs, a cumulative distribution function (CDF) of the RF sizes is obtained (Figure 5A).

The CDF can be differentiated to give the probability density function (PDF) for the RF size distribution (Figure 6). Put another way, the probability of an RF being of a particular size is equal to the slope of the CDF. Sampling these slopes for all line lengths provides the PDF, which represents the probability *density* (probability per unit length) of the RF being that size. Figure 6 shows the numerical estimate of the distribution using the data in Figure 5A. As this experiment was limited to 5 levels of the line length parameter, the resulting graph is coarse. The parameters used in this experimental design were discrete and limited the experimental design’s power due to its fully factorial nature. To more efficiently estimate the maximal pRF size, an algorithm to optimize on the “plateau” boundary as described above complement the analysis (see Yazdanbakhsh & Gori (2008b)).

**Figure 6.**
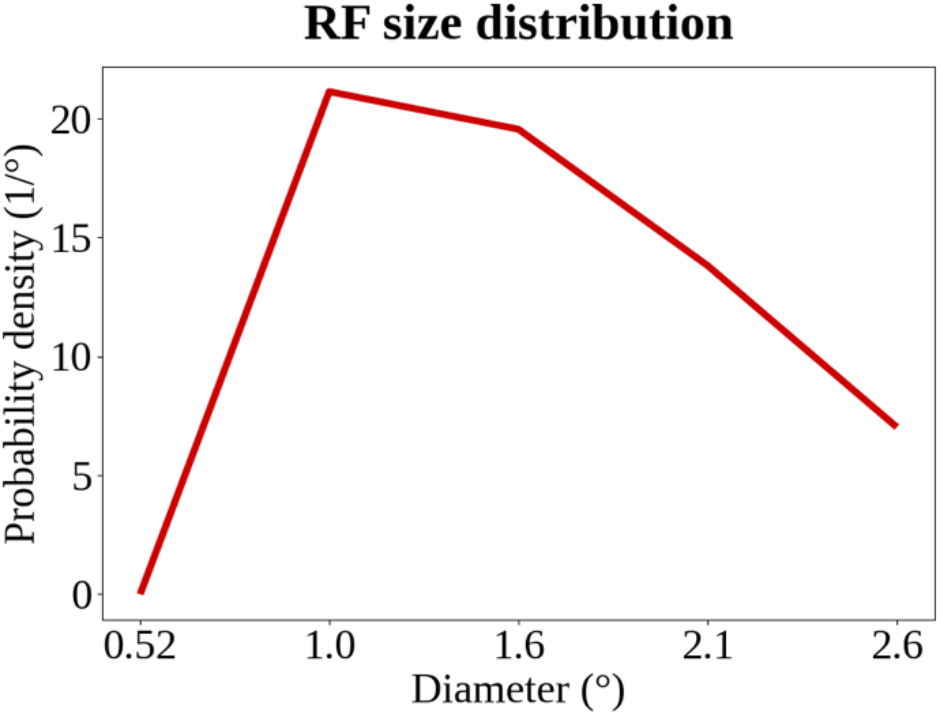
RF size probability density function estimate. This is obtained by finding the relative increases in illusion strength between different values of the line length parameter.

### 4.3 RTLI-based pRF estimation for the evaluation of cognitive abnormalities

We used the novel approach we developed to examine RTLI-related visual processing in conditions associated with changes in pRF size, including autism spectrum disorder (ASD), schizophrenia (SZ), Alzheimer’s disease (AD), and aging.

#### 4.3.1 Autism spectrum disorder

Enlargement of pRF sizes has been described empirically in ASD (Dumoulin and Knapen, 2018; Schwarzkopf et al., 2014). We predict that the observed increase in receptive field sizes, likely decreases the strength of the illusion for individuals with ASD (Figure 7). Specifically, we predict altered illusion strength-line length and illusion strength-stimulus radius relationships in individuals with ASD (Figure 7). As line length increases, the larger RFs observed in ASD imply that a smaller proportion of neurons will be affected by the illusion, decreasing its strength (Figure 7A). A similar effect is theorized with regards to stimulus radius; at all eccentricities, there would be a weaker illusion due to relatively larger RFs (Figure 7B).

**Figure 7.**
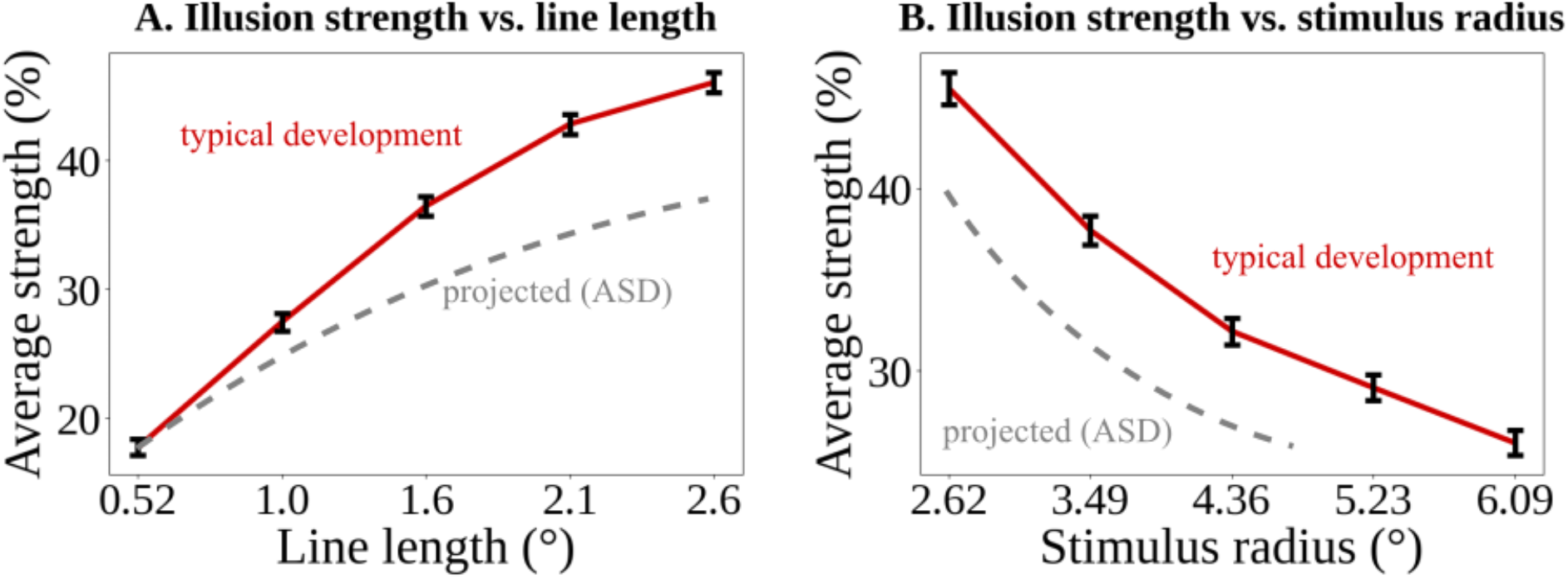
Projected effects of autism spectrum disorder on illusion strength with regards to line length and stimulus radius. A. We predict a decreased illusion strength in individuals with ASD, where divergence from typical development increases at longer line lengths, and B. at all stimulus radii.

#### 4.3.2 Schizophrenia

Patients with SZ exhibit smaller pRFs in several cortical visual areas at all eccentricities (Anderson et al., 2017). Consequently, we predict distinct patterns in both the line length-illusion strength and stimulus radius-illusion strength relationships (Figure 8). As line length increases, the smaller pRFs in SZ are expected to yield a higher percentage of RFs ‘fooled’ by the aperture problem and produce a stronger illusory rotation (Figure 8A). A similar effect is theorized with regards to stimulus radius: at all eccentricities, smaller RFs should lead to a stronger illusion (Figure 8B).

**Figure 8.**
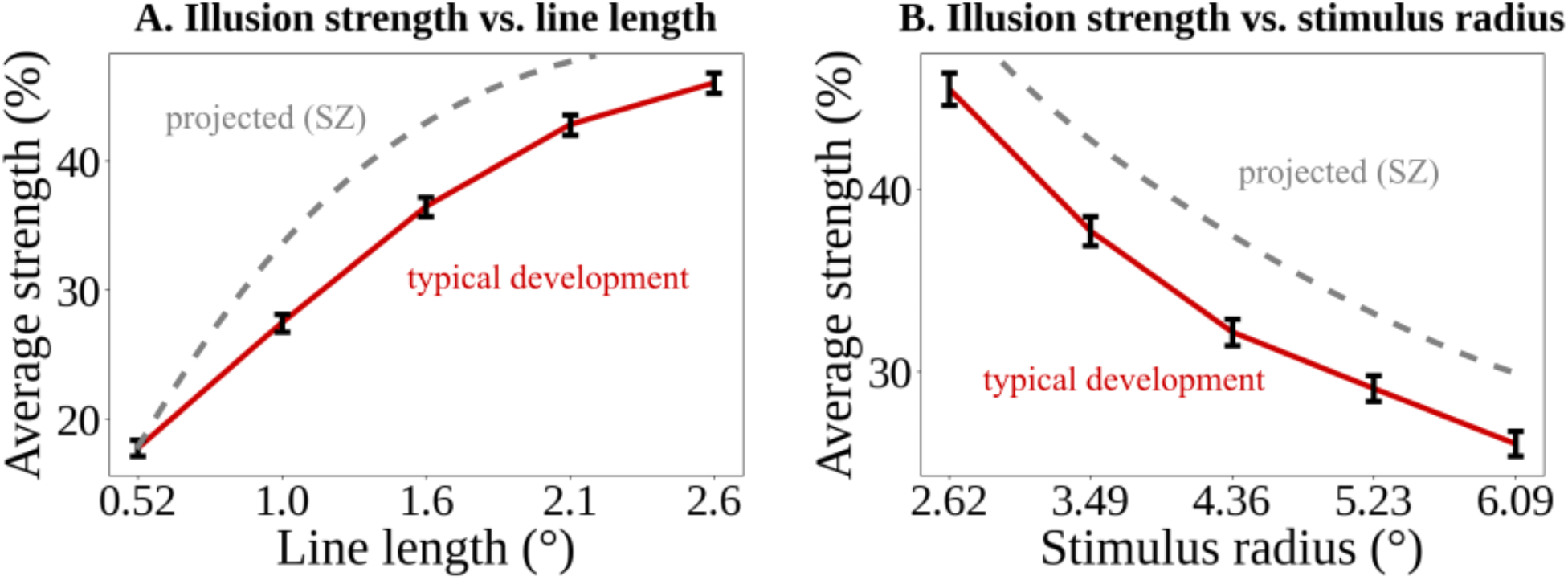
Projected effects of schizophrenia on illusion strength vs. line length and stimulus radius. A. Due to overall smaller pRF sizes in SZ, at each line length, the total percentage of receptive fields affected by the aperture problem are higher than the total percentage of affected receptive fields in typical development leading to higher projected illusion strength. B. Similarly due to smaller pRF at all eccentricities in SZ, we predict increased illusion strength in radius-illusion strength relationship.

#### 4.3.3 Aging and Alzheimer’s disease

Although pRF characteristics remain poorly understood in AD and its prodromal stages, some evidence suggests that aging itself is associated with increases in pRF size, but such deviations due to age were less pronounced as one went up the visual hierarchy. (Brewer & Barton (2014); Silva et al. (2021); Dumoulin and Wandell (2008)). Therefore, in aging populations, we expect a smaller proportion of RFs to be affected by the RTLI illusion due to increased pRF size (Figure 9A). Furthermore, Illusion strength is expected to diverge more from typically developing patterns at smaller radii, with deviations diminishing as eccentricity increases (Figure 9B).

**Figure 9.**
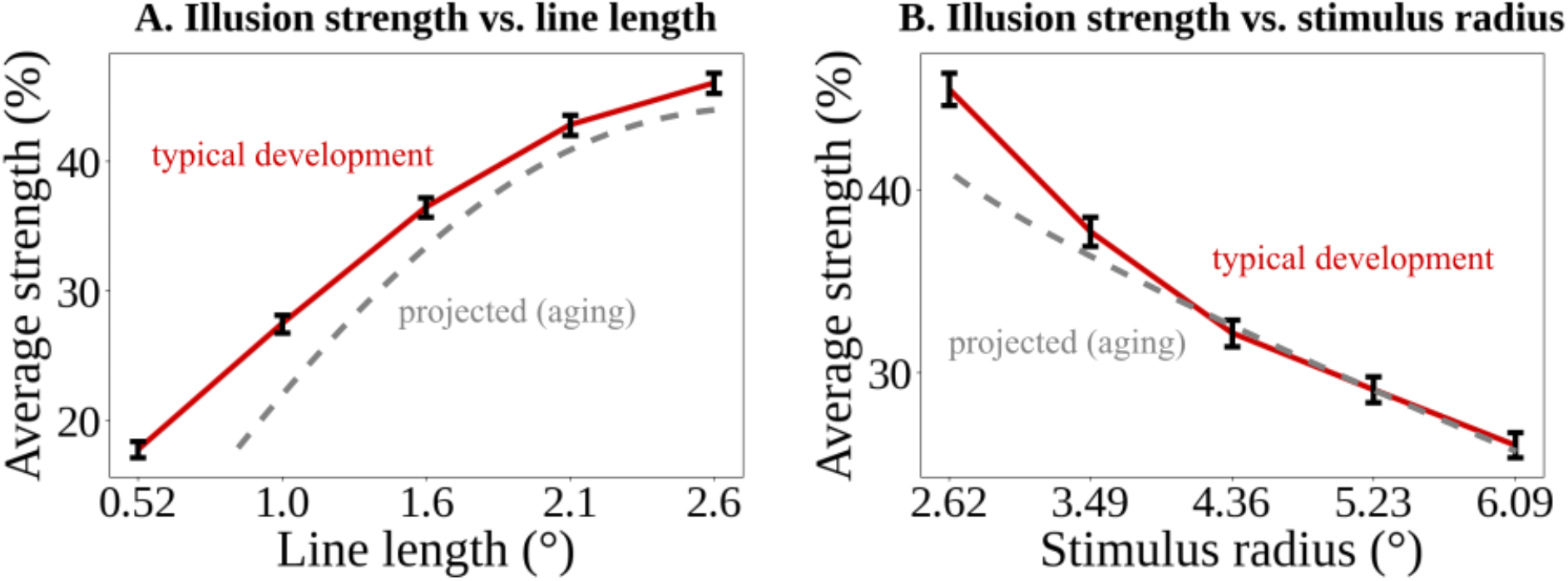
Projected effects of aging on illusion strength with regards to stimulus radius. A. We predict lower illusion strength in aging subjects, with larger discrepancies at lower line lengths. B. We predict lower illusion strength in aging subjects, with larger discrepancies at lower radii and convergence with typical development at higher radii.

## 5. Discussion

The Rotating Tilted Lines Illusion (RTLI) is an optical illusion which appears to rotate as it expands and contracts within the visual field. This illusory rotational effect is caused by the aperture problem (Wallach, 1935) faced by the small receptive fields of V1 (Gori and Yazdanbakhsh, 2008; Yazdanbakhsh and Gori, 2008b). Our study quantifies how illusion strength varies in relation to line length, stimulus radius, and animation speed. Through systematic variation of these parameters, the RTLI test can estimate population receptive field (pRF) size. Our method requires only computer equipment, providing a cost-effective and non-invasive supplement to traditional fMRI and electrophysiological methods in estimating pRF sizes.

### 5.1 RTLI parameters and illusion strength

The present study characterizes the RTLI percept in terms of its spatial and temporal parameters. The RTLI strength measured based on factorial changes of three illusion main variables shown in Figures 5A and 5B are consistent with the aperture problem framework (Christopher C Pack and Born, 2001; Sherbakov and Yazdanbakhsh, 2013; Stumpf, 1911). Illusion strength increases monotonically as line length increases (Figure 5A). This pattern is expected: as line length increases, a greater proportion of receptive fields fall susceptible to the aperture problem, increasing illusory rotation and thus illusion strength. Once line length exceeds the size of most receptive fields, further increases produce minimal change in the proportion of affected RFs, causing illusion strength to plateau. In contrast, illusion strength decreases with higher illusion radius (Figure 5B). As reported by Albright & Desimone (1987), RFs vary significantly with regards to their retinotopic location; their average size increases with more eccentricity, or distance from center of vision. As illusion radius increases, the lines are drawn farther from center of vision, and the RFs processing them are larger. Consequently, the illusion is weakened as the proportion of RFs smaller than the lines is decreased. Lastly, illusion strength decreased with longer animation periods (Figure 5C). This trend reflects the visual system’s ability to resolve the aperture problem over time by integrating input from end-stopped neurons, whose activity is suppressed when contours extend beyond the excitatory region of the RF (Pack et al., 2003; Christopher C. Pack and Born, 2001). At shorter animation periods (0.25–0.5 s), stronger illusory motion was observed, likely reflecting unresolved aperture problems. As animation period increased (i.e., animation speed decreased), illusion strength decreased monotonically, indicating that the visual system had more time to integrate motion signals and infer true motion direction (Christopher C Pack and Born, 2001; Sherbakov and Yazdanbakhsh, 2013). Because this integrative process for all three main parameters of the illusion, including line length, eccentricity, and period, depends on interactions between receptive fields center and their surround, disruption in ceneter/surround mechanisms and/or imbalances between excitation and inhibition may modify, delay, or impair aperture problem resolution. Examining how illusion strength varies with the changes in its three mentioned main parameters in clinical populations may provide insight into spatial and temporal integration processes affected by altered neural dynamics.

### 5.2 pRF estimation for the evaluation of cognitive abnormalities

In our prior work we showed that RTLI-estimated pRF values can be used to distinguish between children with Developmental Dyslexia (DD) and age-matched controls (Gori et al. (2015)). Gori et al. (2015)Here, we used our novel approach to expand the application of RTLI-based pRF estimation and to examine potential correlations with visual processing dysfunction in other conditions associated with changes in pRF size, including autism spectrum disorder (ASD), schizophrenia (SZ), Alzheimer’s disease (AD), and aging.

#### 5.2.1 Autism spectrum disorder

Atypical sensory experiences, affecting up to 90% of individuals with ASD, are a hallmark of the condition and impact every sensory modality, including vision (Tomchek and Dunn, 2007). One consequence of altered visual processing in ASD is reduced susceptibility to several visual illusions, such as the Poggendorff and Zöllner illusions (Poggendorff & Zöllner 1860). Using a computational model, Park et al. (2022) found that reduced susceptibility to the Poggendorff and Zöllner illusions is correlated with an excitation/inhibition imbalance and reduced top-down modulation from higher-order visual areas. These findings align with the MEG and psychophysical studies showing the excitation/inhibition imbalance and reduced feedback connectivity in ASD (Foss-Feig et al., 2013; Seymour et al., 2019).

Cortical feedback from higher-area neurons is inherently non-linear relative to stimulus driven cortical feedforward signals; decreased functional feedback connectivity in autism suggest a decreased non-linearity accompanied by an increased veridical and linear components of the cortical response to input in autism (Khan et al., 2015). Notably, the feedback connections from higher-order visual regions could affect the excitatory and inhibitory signals in V1, thereby modulating the neurons’ receptive field size (de Best et al., 2019). Indeed, larger pRF sizes in ASD have been identified both empirically (Dumoulin and Knapen, 2018; Schwarzkopf et al., 2014) (Schauder et al., 2017) and computationally (Schauder et al., 2017) (Dumoulin and Knapen, 2018; Schwarzkopf et al., 2014). These findings support our prediction that the alterations in receptive field sizes in ASD, either stemming from excitation/inhibition imbalance or from alterations in feedback, likely decrease the strength of the illusion for those with ASD.

While the temporal dynamics of the aperture problem resolution in ASD populations remains relatively uncharacterized, fMRI studies have reported weaker surround suppression in both the MT (Schallmo et al., 2020) and V1 (Flevaris and Murray, 2015). The underlying causes of these alterations are unclear but are thought to involve a combination of feedforward, lateral, and feedback mechanisms. Although current evidence is insufficient to form a concrete prediction, reduced suppression suggests that the inhibitory processes underlying end-stopping may be less effective in ASD, potentially leading to slower or less efficient resolution of the aperture problem. This hypothesis could be tested by examining the relationship between animation period and illusion strength, where one might expect weaker time-dependence or slower convergence toward the true motion percept in ASD populations.

#### 5.2.2 Schizophrenia

Schizophrenia has been described by scientists, beginning in the 1950s, as featuring deficits in visual perceptual organization among its most significant characteristics (Silverstein and Keane, 2011). Dysfunction of visual information processing was assessed with visual masking tasks and found to occur before and after the onset of frank psychosis, suggesting that visual deficits may represent a stable marker for SZ (Perez et al., 2012). People with SZ experience several perceptual abnormalities, including aberrations in contrast sensitivity (Dugan et al., 2024; Slaghuis, 1998; Zhu et al., 2023), poor orientation discrimination (Tibber et al., 2015), and impaired motion processing (Chen et al., 2004; Kim et al., 2006; Zhu et al., 2023). Further, center-surround suppression for motion is abnormally weaker in SZ patients, particularly those with severe negative symptoms, as compared to control groups (Tadin et al., 2006).

The reduced surround suppression mechanism in patients with SZ is thought to be preceded by an imbalance between cortical excitation and inhibition. This abnormality has previously been proposed as the mechanism responsible for altered susceptibility to various illusions (Dakin et al., 2005; Yoon et al., 2009). Heeger Zenger-Landolt & Heeger (2010), using a moving checkboard pattern and fMRI, have localized the impaired surround suppression mechanism to V1, V2 and V3, with the strongest effects in the latter two cortices. Difference-of-gaussian models applied to SZ show size reduction of the central excitatory component of the pRF in V1 and V2, as well as a narrowing and shallowing of the inhibitory surround in V1, V2, and V4 (Anderson et al., 2017).

Moreover, surround suppression has been closely linked to magnocellular (M-cell) neurons. The perceptual differences observed in individuals with SZ (compared to normal) are linked to a semi-selective dysfunction in the magnocellular visual pathway (Butler et al., 2005; Martínez et al., 2008), therefore with a bias toward smaller pRF sizes in SZ patients. The aforementioned factors are consistent with smaller pRFs in V1, V2 and V4 at all eccentricities (Anderson et al., 2017). In line with this data, our approach shows that smaller pRFs in SZ lead to a stronger illusion percept (Figure 8). Information on temporal processing dynamics that could inform predictions of animation period--illusion strength relationships in SZ populations remains limited. The theorized reduction in surround suppression may similarly contribute to slower or impaired resolution of the aperture problem—for example, manifesting as a more gradual decline in illusion strength with increasing animation period.

#### 5.2.3 Aging and Alzheimer’s disease

Alzheimer’s Disease (AD) presents another facet of cognitive impairment in which the RTLI task warrants exploration. Because visual deficits manifest as early symptoms of AD, alterations within the visual cortex may precede the clinical onset, offering a possible avenue for early detection. Notably, AD is not a singular, homogeneous disease but rather an end-point presentation (Emery, 2011). Trajectories leading to AD typically fall under two categories: (1) normal aging progression to AD and (2) depressed elderly progression to AD. Although pRF characteristics remain poorly understood in AD and its prodromal stages, some evidence suggests that aging itself is associated with changes in pRF size.

Brewer & Barton (2014) used pRF size analysis ascertained by computational methods, compared against fMRI in Dumoulin and Wandell (2008), to create a model for pRF size mapping. A MAVONA analysis by Brewer and Barton revealed significant increase in pRF size in the central 3° for aging participants (ages 57-70), but not in the peripheral 3-10° of V1, V2 and hV4. In V3, there was a marginally significant increase in pRF size in both central 3° and peripheral 3-10°. Silva et al. (2021) similarly found (using fMRI) that aging is associated with larger pRF size in V1, V2, and V3, but such deviations due to age were less pronounced as one went up the visual hierarchy. Therefore, this is in line with our prediction that in aging populations a smaller proportion of RFs will be affected by the RTLI illusion due to increased pRF size, and this divergence will be more prominent at smaller radii, with deviations diminishing as eccentricity increases.

In relation to the AD, Brewer and Barton (2014) conducted a similar analysis in two subjects with mild Alzheimer’s disease. They observed irregularities in visual field map size and organization, which differed between the two AD subjects. However, both showed similar reductions in pRF sizes in peripheral regions of V2, V3 and hV4. Prediction trends for AD are not presented here, as current literature and subject pools remain limited on pRF size deviations in this population.

Pathological hallmarks of AD are typically measured through neuroimaging, cerebrospinal fluid analysis, or diagnosed post-mortem (Javaid et al., 2016). As Brewer and Barton’s findings suggest, alterations in pRF size along the pre-AD trajectory may serve as early indicators of disease onset.

Unlike the three conventional diagnostic methods, the RTLI task is non-invasive, cost-effective, and scalable to larger sample sizes, making it a promising tool for assessing pRF size changes across early, moderate and late stages of AD.

## 5. Conclusions

In the RTLI, a circle formed by tilted lines seemingly appears to rotate when it expands or contracts in the visual field. We present a computer-based mechanism for animating and measuring the strength of the RTLI. Results are shown for measuring illusion strength against three parameters: line length, stimulus radius and animation period. Using the length at which illusion strength started to plateau as an estimate of the maximum pRF size, we estimated pRF size of between 1.56 and 2.09° for healthy subjects. We further present a pRF size distribution obtained from altering the line length parameters.

Moreover, as a cost-effective and non-invasive method of estimating pRF sizes, the RTLI illusion is a pRF size estimation method that can be used along other methods for early diagnosis of several disorders which implicate visual impairments, including autism spectrum disorder, schizophrenia and Alzheimer’s disease. Using results obtained by neuroimaging and computational methods previously obtained in the field, we present predicted trends of illusion strength vs. line length and stimulus parameters for these groups. Future endeavors may include using this approach to study clinical populations with disorders beyond those discussed in this study, including patients with damage to visual areas in the brain due to stroke, traumatic head injury, chronic encephalopathy, or patients treated with hemispherectomy for other conditions.

## Acknowledgements

We thank the members of the Computational Neuroscience and Vision Laboratory for their participation and valuable feedback during the development and conducting of experiments.

## CRediT authorship contribution statement

**Chris Bao:** Conceptualization, Software, Validation, Formal Analysis, Investigation, Data Curation, Writing – original draft, Visualization. **Emma Martin:** Investigation, Data Curation, Writing – original draft, Visualization, Project Administration, Writing – Review and Editing. **Basilis Zikopoulos:** Conceptualization, Writing – review & editing, validation. **Arash Yazdanbakhsh:** Conceptualization, Methodology, Software, Validation, Formal Analysis, Resources, Data Curation, Writing – original draft, Writing – Review and Editing, Supervision, Project Administration.

## Supplementary materials

## Appendix A. Experiment files

The experiment code and previous versions are available online at https://github.com/ChrisBaoCC/RTLIExperiment

## Appendix B. Aperture problem and RTLI lines orientation

The illusion arises from the component of motion perpendicular to the line segment’s orientation (shown in blue), which itself contains a component on the tangential direction (shown in red).

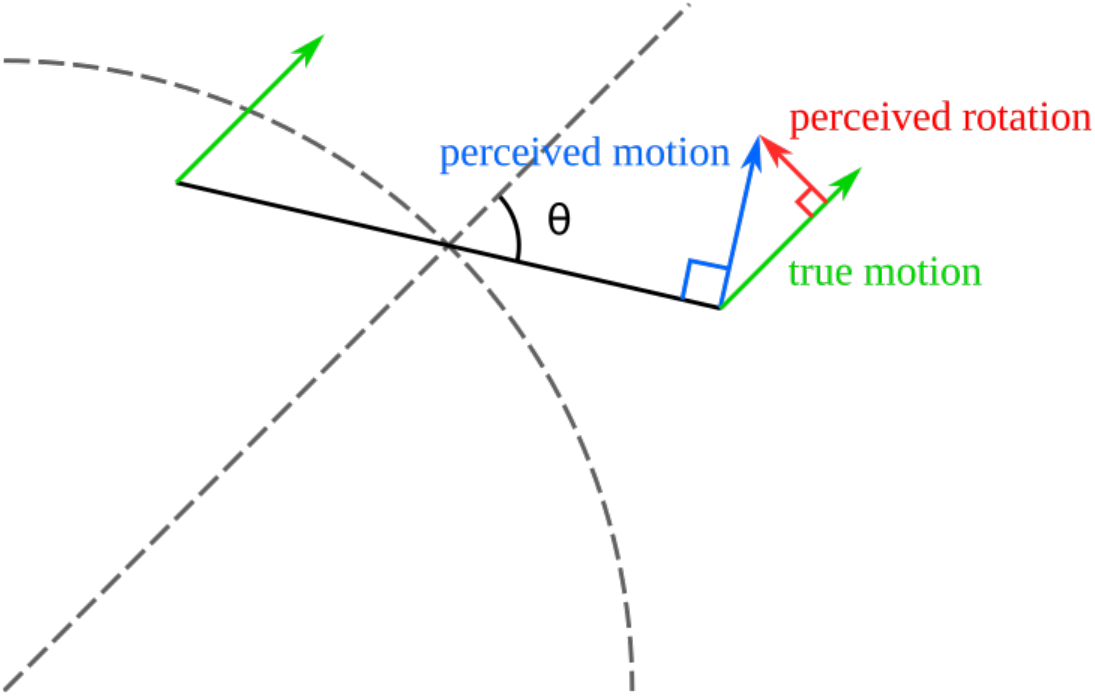

We define *θ* as the angle between the radial line and the line segment and *v* to be the true velocity (which is completely radial, shown in green). Due to the aperture problem, the perceived direction of motion is the component of *v* perpendicular to the line, or *v* sin(*θ*). The speed of the perceived rotation is equal to the tangential component of perceived motion, which equals *v* cos(*θ*) sin(*θ*). Theoretically, the value of *θ* that maximizes this function is π/4 radians or 45°.

## Appendix C. Violin plots

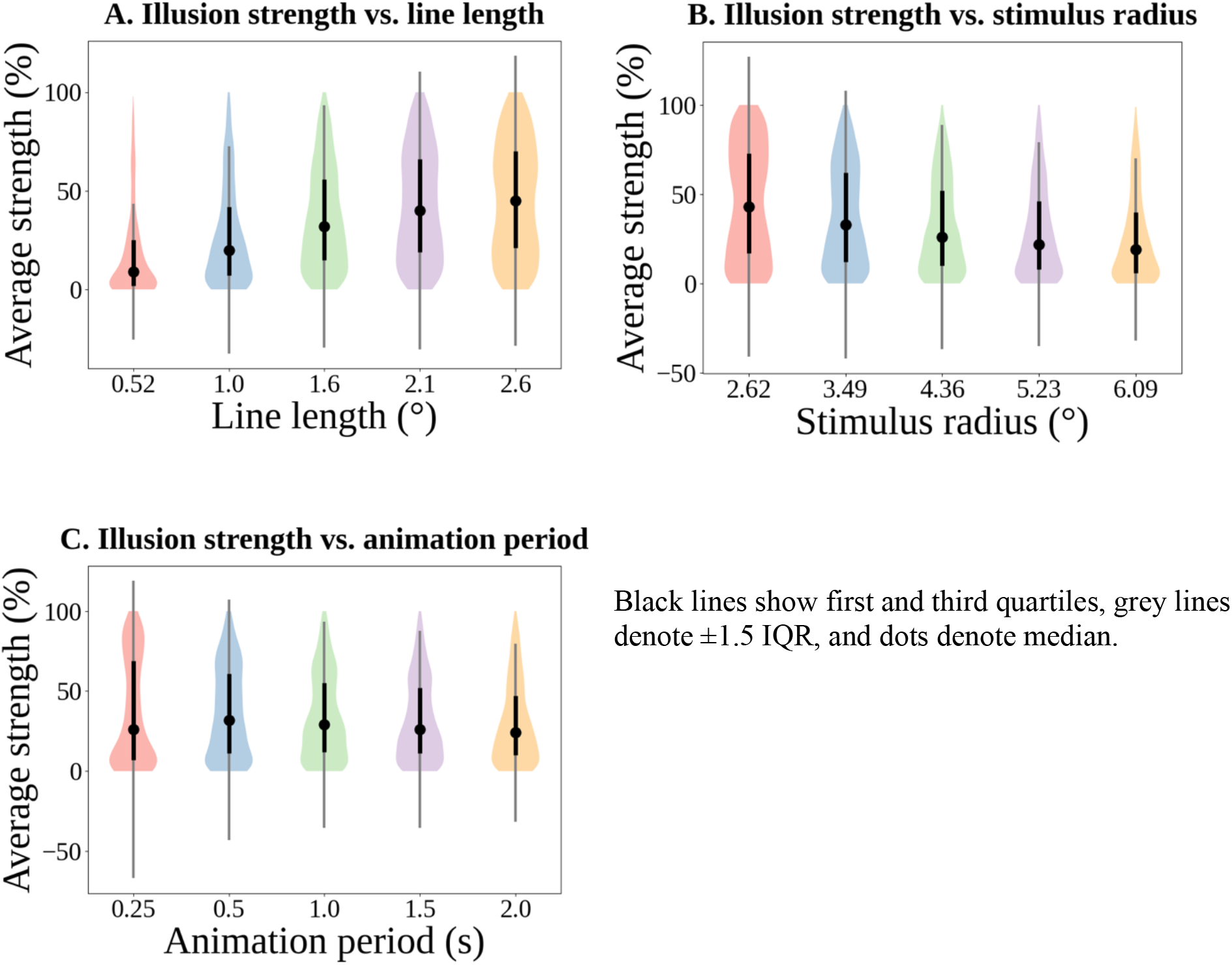

Black lines show first and third quartiles, grey lines denote ±1.5 IQR, and dots denote median.

## Appendix D. Power analysis

Additional analysis was conducted post-hoc to determine the power of this experiment using the PowerSampleSize software by Dupont & Plummer, which may be found at https://biostat.app.vumc.org/wiki/Main/PowerSampleSize. Logged results for linear regressions are as follows.

### Illusion strength vs. line length

We are planning a study with 6125 subjects × trials evenly divided into 5 groups receiving xvar at levels 0.524, 1.05, 1.57, 2.09, and 2.62. Prior data indicate that the standard deviation of the regression errors will be 0.74. If the true slope of the line obtained by regressing yvar against xvar is 0.2399, we will be able to reject the null hypothesis that this slope equals zero with probability (power) 1.000. The Type I error probability associated with this test of this null hypothesis is < 0.05.

### Illusion strength vs. stimulus radius

We are planning a study with 6125 subjects × trials evenly divided into 5 groups receiving xvar at levels 150, 200, 250, 300, and 350. Prior data indicate that the standard deviation of the regression errors will be 70.711. If the true slope of the line obtained by regressing yvar against xvar is −0.0954, we will be able to reject the null hypothesis that this slope equals zero with probability (power) 1.000. The Type I error probability associated with this test of this null hypothesis is < 0.05.

### Illusion strength vs. animation period

We are planning a study with 6125 subjects × trials evenly divided into 5 groups receiving xvar at levels 25, 50, 100, 150, and 200. Prior data indicate that the standard deviation of the regression errors will be 64.031. If the true slope of the line obtained by regressing yvar against xvar is 0.0425, we will be able to reject the null hypothesis that this slope equals zero with probability (power) 1.000. The Type I error probability associated with this test of this null hypothesis is < 0.05.

